# The nucleus forms a dynamic contact with the plasma membrane to maintain the glandular epithelial architecture

**DOI:** 10.1101/2025.06.13.659552

**Authors:** Megane Rayer, Keeley Mui, Kavya Pendyala, Johnny Rockenbach, Susumu Antoku, Tanmay P. Lele, Gregg G. Gundersen

## Abstract

The maintenance of epithelial architecture relies on precise mechanical and biochemical cues. Recent studies reveal an unexpected role for the nucleus in maintaining epithelial architecture, but how the nucleus is physically and molecularly integrated into epithelia remains unclear. Here, we identify a dynamic basal actin spot that links the nucleus to plasma membrane β1-integrin through the linker of nucleoskeleton and cytoskeleton (LINC) in 3D breast acini. Depletion of LINC complex nesprin-2G, SUNs or FHOD1 disrupts nuclear positioning and inhibits lumen formation. Activation of a nesprin-2 degron causes acute loss of the basal actin spot and collapse of acini. Active Src and β1-integrin accumulate in the basal actin spot and Src activity is required to prevent collapse of acinar structure. These findings reveal an unexpected mode of nuclear-plasma membrane contact that we propose homeostatically regulates intracellular contractility through a Src signaling pathway to maintain global epithelial architecture.

## Introduction

The architecture of tissues and organs are essential to their function, and defects in their development or maintenance can lead to disease. Mechanical properties are fundamental for shaping and preserving tissue and organ architecture. The nucleus is the largest and most rigid organelle in cells and this makes it a hub of mechanotransduction. At the center of this hub, is the linker of nucleoskeleton and cytoskeleton (LINC) complex comprised of KASH proteins (nesprins in vertebrates) and SUN proteins in the outer and inner nuclear membranes, respectively. These proteins associate in the perinuclear space to form a physical connection from the cytoskeleton to the nuclear lamina and nucleoplasm (*1*). The forces transmitted by the LINC complex can move and position the nucleus(*2–5*), contribute to nuclear shape (*6, 7*), alter nuclear pore function (*8*); and affect chromatin organiztion and gene expression (*9–11*). Forces impinging on the LINC complex can also affecting signaling both within the nucleus (*12*) and in the cytoplasm (*13, 14*). How these various activites are coordinated to build tissue and organ architecture is unknown.

In the original phenotypic screens in model organisms that identified LINC complex components, striated muscle, nerve and epithelia tissues were affected (*15*). Since these early studies, we have developed a keen understanding of how the LINC complex contributes to muscle and nervous tissue (*3, 16–18*). Mechanistic studies on epithelia have lagged. Progress has been made to understand the role of the LINC complex in the stratified epithelia of skin, where the LINC complex plays a key role in early differentiation and in homeostatic wound heading (*10, 11*). More recent studies with epithelial 3D organoids or acini have highlighted the importance of the nucleus and the associated LINC complex in maintaining polarized epithelial architecture. In acini generated from MDCK kidney cells or MCF10A breast cells, disruption of the LINC complex leads to increased contractility, lumenal collapse and loss of overall epithelial organization, (*19*). Over time, cells in the collapsed acini move outward (evert) generating clusters of cells with inverted apical-basal polarity (*20*). Other factors that lead to acinar collapse and eversion include elevating Rho GTPase contractility and disrupting β1-integrin (*20*).

Most cells express more than one nesprin and SUN protein. Glandular epithelia express four (nesprins 1-4) and both of the widely expressed SUN proteins (SUN1 and SUN2) (*10, 11*). The dominant negative constructs used in the previous studies of the LINC complex in epithelial acini disrupt all of these, so it is unclear whether acinar architecture requires a specific nesprin-SUN protein complex, whether multiple types of LINC complexes are involved, or whether specific nucleocytoskeletal connections are required. Addressing this question has relevance beyond basic organ biology. In individual carcinoma cases, individual rather than global LINC complex species are downregulated. For example, in breast cancer, one of the LINC complex proteins (nesprin-2, SUN1 or SUN-2) or the associated lamin A/C was downregulated in virtually every individual case examined (*21–24*). As altered lumenal structure, cell polarity and contractility are hallmarks of cancer (*25–27*), understanding whether changes in the LINC complex alter these features may lead to further appreciation of the contribution of mechanical factors in cancer.

We report here that a specific set of LINC complex components including SUN2, nesprin-2G (G refers to the giant isoform) and the nesprin-2G associated formin FHOD1, is required to form normal breast acinar architecture and that it is the actin functionality of nesprin-2G (and FHOD1) that is required. Searching for a nuclear-associated actin filaments, we identify a basal spot of F-actin in close association with the nucleus. This spot forms transiently, anchors the nucleus and using a degron of nesprin-2, is required to maintain acinar architecture. Based on colocalization of plasma membrane β-1 integrin and SUN2 with the actin spot, we propose that the structure represents a novel nuclear-plasma membrane contact site. As active Src colocalizes with the contact site and Src inhibitors also disrupt acinar architecture, we propose that the nuclear-plasma membrane contact site represents a signaling center that regulates cellular contractility to maintain acinar structure. We propose the name: <u>P</u>lasma membrane and <u>A</u>ctin-associated <u>N</u>uclear Inhibitor of <u>C</u>ontractility (PANIC) button.

## Results

### Specific LINC complex proteins and functionality are required for lumen formation

Previous studies implicating the LINC complex in glandular epithelia architecture employed dominant negative constructs that indiscriminately disrupt all LINC complexes (*19*). To identify specific LINC complexes responsible for acinar formation, we conducted a mini-screen of LINC complex proteins in non-transformed human MCF10A breast epithelial cells (*25*) and normal (non-transformed) murine mammary gland cell (NMUMG). Knockdown of nesprin-2, but not nesprin-1 and either SUN1 or SUN2 prevented formation of MCF10A acini with hollow lumens (Figure 1a,b; Supplementary Fig. 1a). Similar results with nesprin-2G knockdown were obtained in NMuMG cell acini (Figure 1c,d and Supplementary Fig 1b). Thus, specific LINC complex components are required for lumen formation in both human and mouse breast epithelial cells.

**Figure 1.**
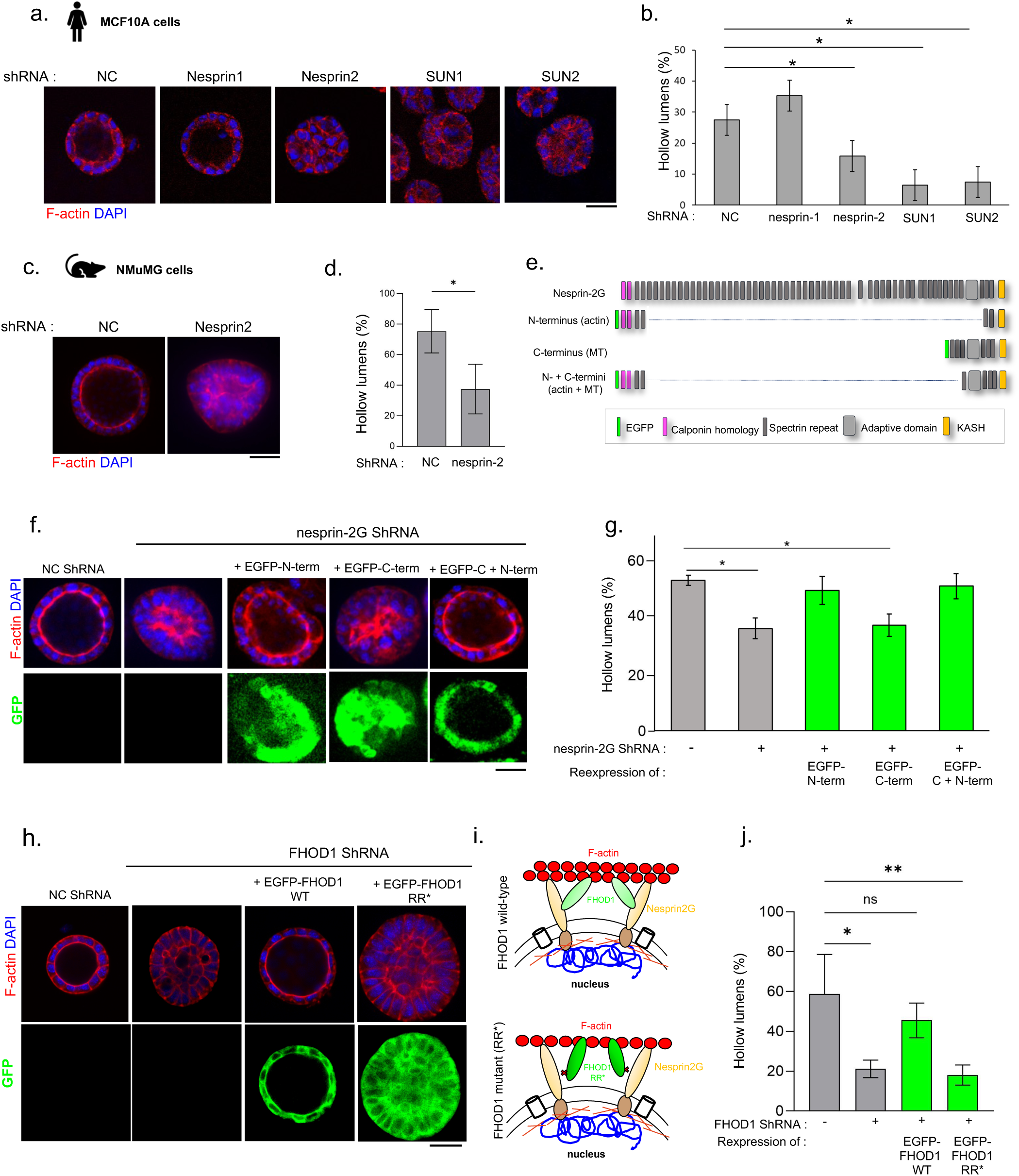
Specific LINC components are required for acini lumen formation. **a)** Representative images of fixed acini formed by MCF-10A cell lines expressing shRNAs targeting individual LINC complex gene. NC, noncoding control. Acini were stained with phalloidin (F-actin, red) and DAPI (DNA, blue). **b)** Histogram showing the quantification of the percentage of MCF10A acini with hollow lumens. Statistical test: One-way Anova with post hoc Tukey: * p ≤ 0.05 Data from 3 independent experiments: n(NC) = 212 acini, n(Nesprin-1) = 300 acini, n(Nesprin-2) = 280 acini, n(SUN1) = 171 acini, n(SUN2) = 221 acini. **c)** Representative images of acini formed by NMuMG cell lines after knockdown of nesprin-2G using shRNA compared to noncodling control (NC) shRNA. Acini are stained with phalloidin (F-actin, red) and DAPI (DNA, blue). **d)** Histogram shows the quantification of the percentage of NMuMG acini with hollow lumens after treatment with the indicated shRNAs. Unpaired t-test: * p ≤ 0.05. Data from 3 independent experiments: n(NC) = 124, n(Nesprin-2) = 167. **e)** Schematic representation of full-length nesprin-2G and three fragments (N-terminus interacting with actin, C-terminus interacting with microtubules, N-terminus plus C-terminus interacting with both). All are tagged by EGFP. **f)** Representative images of acini generated by NMuMG acini stablu knocked down for nesprin-2G and re-expressing different EGFP-tagged fragments of nesprin-2G (EGFP positive acini). **g)** Histogram showing the quantification of the percentage of acini with hollow lumens for cells treated as in f. One-way Anova with post hoc Tukey Statistical test: *p* values ≤ 0.05 indicated as * for significance. N = 3 independent experiments; n(NC) = 276 acini, n(nesprin-2 shRNA) = 341acini, n(nesprin-2 shRNA + N-term) = 288 acini, n(nesprin-2 shRNA + C-term) = 333 acini, n(nesprin-2 shRNA + N+C-term) = 306 acini. **h)** Representative images of acini formed by NMuMG cells expressing FHOD1 shRNA against FHOD1, with rescue by either FHOD1 wild-type or the nesprin-2 binding deficient mutant FHOD1 R136A/R137A. Both were EGFP tagged. **i)** Top: Diagram illustrating the interaction between nesprin-2 and FHOD1 to promote interaction with F-actin. Bottom: Diagram showing the FHOD1 R136A/R137A mutant, which is unable to bind nesprin-2G. **j)** Histogram showing the quantification of the percentage of acini with hollow lumens. N = 3 independent experiments; n(NC) = 138 acini, n(FHOD1-4) = 159 acini, n(FHOD1-WT) = 185 acini, n(FHOD1 R136A/R137A) = 168 acini. Statistical significance by One-way Anova with post hoc Tukey test: * *p* values ≤ 0.05; ** *p* values ≤ 0.01. Scale bars, a, c, f, h: 30 µm

Nesprin-2G interacts with both the actin and microtubule cytoskeletons (*18, 28, 29*). It links F-actin through amino-terminal paired calponin homology domains, whereas microtubules interact with its C-terminal region through motor proteins and their adaptors (*30*). To identify which domain is required for normal acinar architecture, we re-expressed amino- and carboxy-fragments of nesprin-2G capable of interacting with actin filaments, microtubules, or both elements in NMuMG cells knocked down for nesprin-2G (Figure 1e). Note that the amino terminal fragment was directed to the nucleus by attaching it to the KASH domain. These constructs have previously been shown to be functional in multiple cell contexts (*5*). Expression of these constructs was confirmed by EGFP fluorescence (Figure 1f). Re-expression of both constructs containing the amino terminal actin-binding domain successfully rescued lumen formation in nesprin-2G-depleted cells (Figure 1f,g). In contrast, re-expression of the carboxy terminal fragment mediating microtubule interaction failed to rescue lumen formation (Figure 1f,g). Thus, actin but not microtubule binding of nesprin-2G is crucial for forming normal acini architecture.

To further test the requirement for nesprin-2G connection to actin filaments, we examined whether the formin FHOD1 was required. FHOD1 is a widely expressed formin that physically interacts with nesprin-2G spectrin repeats located near its amino terminus and is required for many nesprin-2G dependent functions that require actin binding (*31–34*). FHOD1 weakly stimulates actin polymerization but strongly bundles actin filaments contributing to their ability to resist mechanical stress (*31, 32*). Knockdown of FHOD1 in NMuMG cells using an shRNA impaired lumen formation (Figure 1h,j; Supplementary Fig. 1c). Re-expression of wild-type human FHOD1 in FHOD1-depleted NMuMG cells rescued the formation of lumens confirming knockdown specificity (Figure 1h,j). To test whether the interaction between FHOD1 and nesprin-2G was critical for lumen formation, we expressed a FHOD1 mutant (R136A R137A) that disrupts its binding to nesprin-2G (*34*) in FHOD1-depleted NMuMG cells (Figure 1i). Cells expressing this mutant were unable to restore lumen formation (Figure 1h,j). Combined with the data from nesprin-2G fragment expression, these data strongly support the idea that nesprin-2G’s actin binding functionality is necessary for acinar lumen formation.

### Conditional nesprin-2G degradation disrupts acini lumen maintenance

Previous work showed that an inducible dominant negative construct caused collapse of acini lumens (*19, 20*). To test whether nesprin-2G also plays a role in lumen maintenance, we developed a conditional degradation system targeting nesprin-2G based on the AID2 degron system (*35, 36*) (Figure 2a). First, we generated NMuMG cells stably knocked down for nesprin-2G using shRNA. In a second step, we transfected the cells with a mAID-EGFP-mN2G (mN2G-degron) construct that contains the auxin-inducible degron tag, a EGFP tag and the actin binding functionality of nesprin-2G, and in a third step, with the degradation enzyme OsTIR1. We confirmed that the endogenous nesprin-2G protein was effectively silenced by the shRNA and that miniN2G-degron was expressed, localized properly to the nucleus and rescued lumen formation in knockdown cells (Figure 2b-d). We then validated the degron system in NMuMG cell cultures where the miniN2G-degron signal disappeared within 30 min of activating the degron with the auxin 5-Ph-IAA as assessed by both immunofluorescence staining and western blotting (Figure 2b,c). To assess whether activating the mN2G-2 degron affected lumen maintenance, we added 5-Ph-IAA after seven days of acini development when lumens had properly formed. In fixed cells, we observed loss of the mN2G-degron levels and reduced numbers of acini with hollow lumens (Figure 2d,e). In live cell movies, we observed lumen collapse after exposure of mN2G-degron cells to auxin (10 of 10 movies) compared to no collapse of mN2G-degron acini to exposed DMSO vehicle (6 of 6 movies) (Figure 2f) Addition of 5-Ph-IAA had no effect on lumen formation in wild-type NMuMG acini (Supplementary Fig. 2b). Together, these results demonstrate that N2G is essential not only for lumen formation but also for its maintenance.

**Figure 2.**
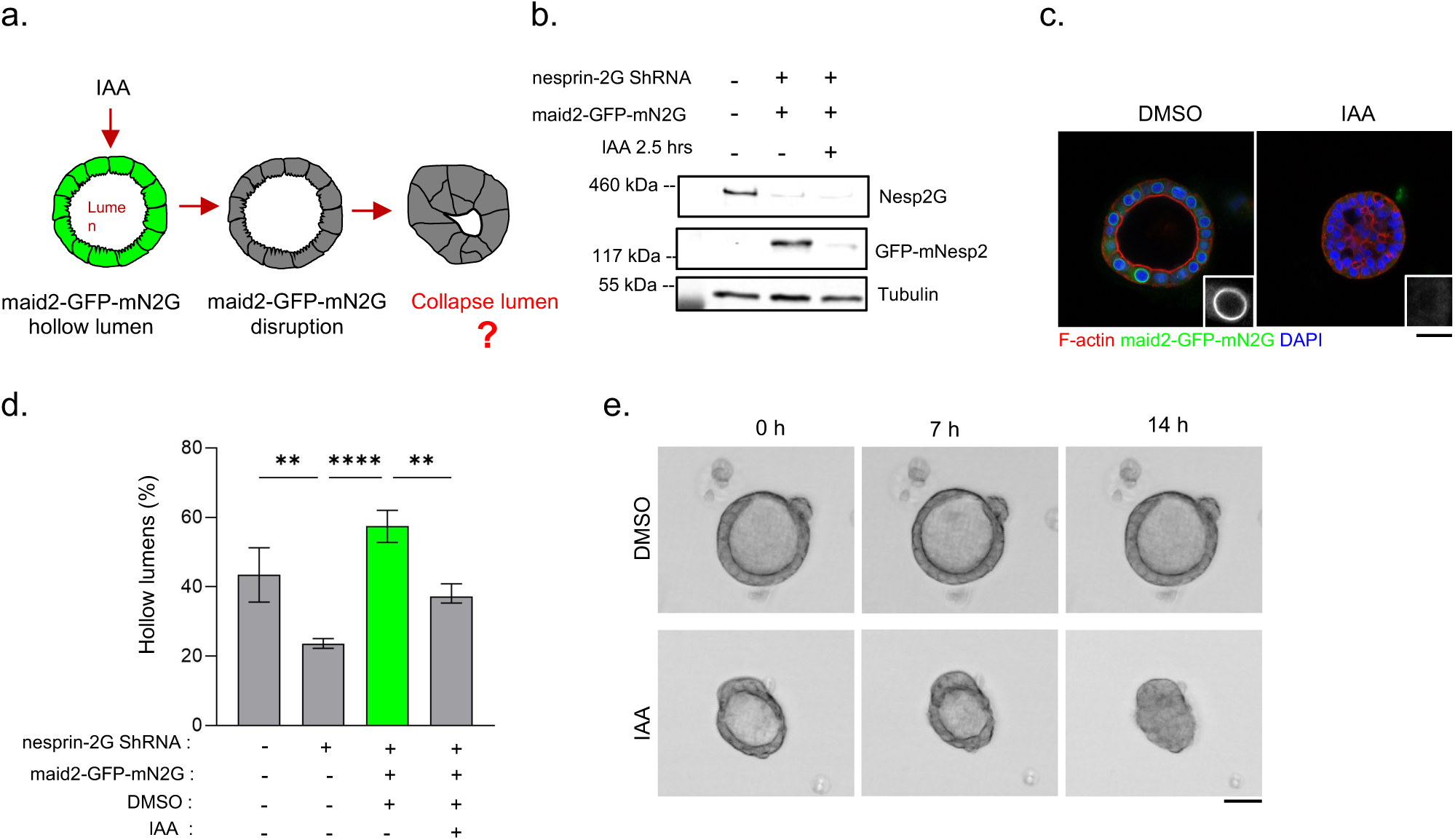
Conditional degradation of nesprin-2G disrupts maintenance of acini lumens. **a.** Diagram illustrating the degron strategy in which addition of 5-Ph-IAA (“IAA”) induces the degradation of mAID-EGFP-mN2G in NMuMG acini stably knockdown for nesprin-2G. The question is whether loss of the mN2G degron will result in collapse of acini. **b.** Western blot showing lane 1, shRNA depletion of endogenous nesprin-2G (Nesp2G), lane 2, expression of the mN2G degron (EGFP-mNesp2) and lane 3, loss of the mN2G-degron after 2.5 h treatment with IAA. Tubulin is a loading control. **c.** Representative images showing, on the left, a hollow lumen in DMSO-treated acini with nuclear mAID-EGFP-mN2G localization, and on the right, a collapsed lumen after 24 h treatment with 5-Ph-IAA, accompanied by the loss of the mAID-EGFP-mN2G signal. Cells were stained for F-actin (phalloidin, EGFP and nuclei (DAPI). **d.** Histogram showing the proportion of acini with hollow lumens in the mN2G degron system. Shown are results for NMuMG cells knocked down for nesprin-2G with shRNA, and nesprin-2G knockdown cells reexpressing the mN2G degron (mAID2-EGFP-N2G) treated with DMSO or 5-Ph-IAA. Note rescue of acini lumens with the mN2G degron and subsequent loss after induce the degradation of the mN2G degron with IAA. Data from at least 3 independent experiments. n(NC)= 146 acini, n(N2) = 240 acini, n(DMSO) = 178 acini, n(IAA) = 148 acini. Statistical test: One-way Anova with post hoc Tukey: ** *p* value ≤ 0.01, **** *p* value ≤ 0.0001. **e.** Bright field time-lapse imaging of NMuMG acini knocked down for nesprin-2G and expressing the mN2G degron. Movies show the maintenance of a hollow lumen in vehicle control (DMSO) compared to lumen collapse (at 14 h) in response to 5-Ph-IAA. Similar results were observed in 9 (DMSO) and 11 (IAA) movies. Scale bars c, e: 30 µm

### A dynamic basal actin accumulation contacts and anchors the nucleus

The data from our knockdown and rescue experiments (Figure 1f-g) suggested there may be a cytoplasmic actin array associated with the nucleus via nesprin-2G that was important for maintaining aciniar architecture. To search for this, we stained cells for F-actin and nuclei and looked for actin arrays in the proximity of the nucleus. As expected, we observed F-actin in the apical domain and the cell-cell junctions and that these arrays rarely overlapped with the nucleus (Figure 3a). In contrast, we observed a basal actin “spot” in close association with the nucleus in a subset (∼30%) of the cells. In over 50% of the cases when the actin spot was present, the basal aspect of the nucleus adjacent to the actin spot was indented altering nuclear circularity (Figure 3b-c). Super-resolution confocal imaging revealed that the actin spot was localized toward the center of the basal surface and was not connected to the cortical ring of junctional actin (Figure 3d). Similar appearing basal actin puncta have been previously observed by others in MDCK cells, although the correlation with nuclear proximity and shape was not described (*37*).

**Figure 3.**
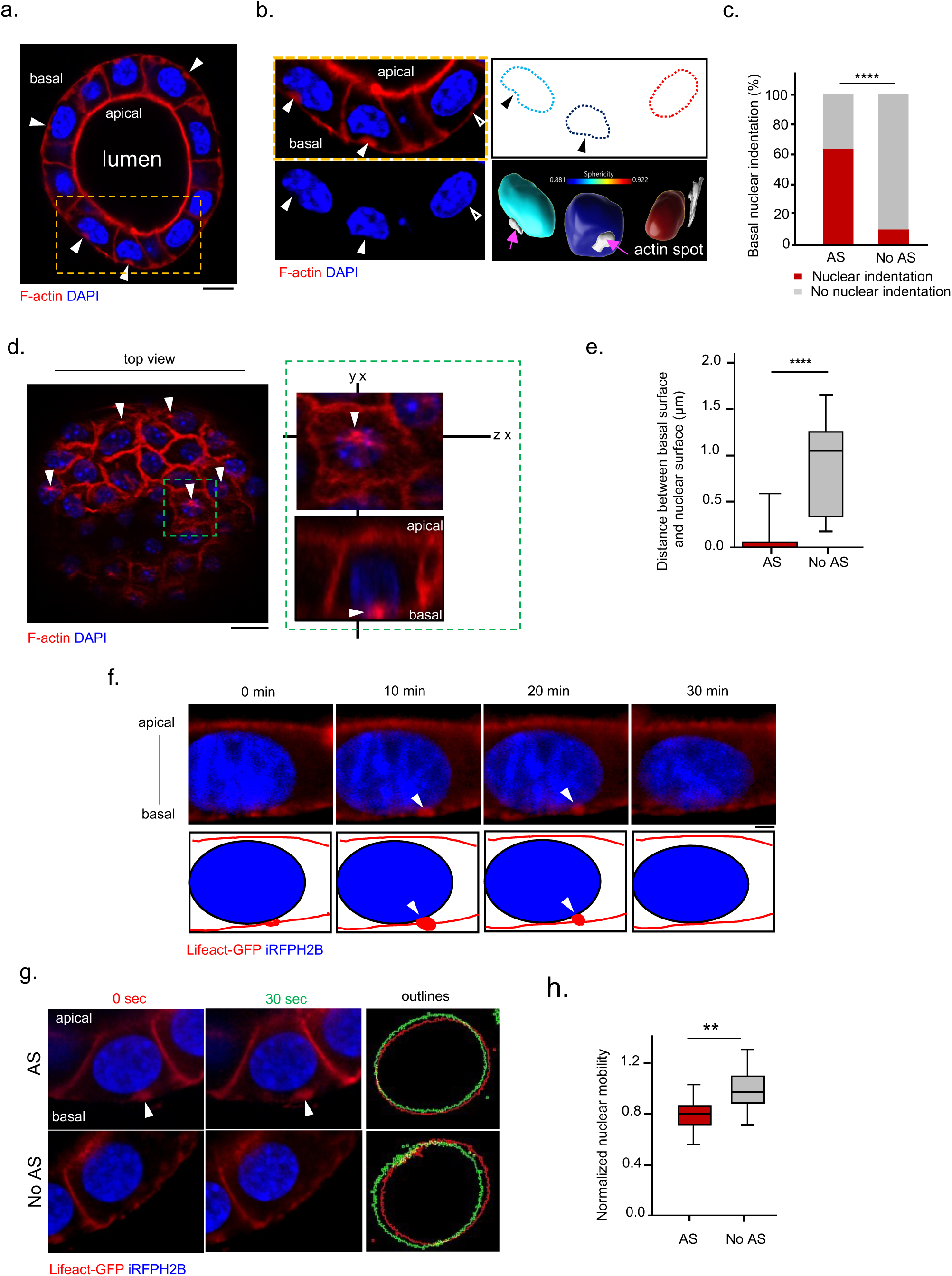
A basal actin spot forms adjacent to the nucleus. **a.** Representative confocal image of an acini showing F-actin (phalloidin) organization relative to the nucleus (DAPI). White arrowheads indicate basal actin spots. The yellow dotted box highlights three nuclei, shown at higher magnification in panel b. Scale bar, 10 µm. **b.** Magnified view of the three nuclei from panel a. The two nuclei in close proximity to actin spots are indented (filled arrowheads). The third nucleus, not in association with an actin spot, remains spherical (empty arrowheads). Bottom right: 3D surface reconstruction of nuclei using IMARIS with color-coded sphericity. Pink arrow indicates the actin spot contact. The two nuclei in close apposition with an actin spot show reduced sphericity. **c.** Quantification of nuclei exhibiting indentation in relation to the presence of an adjacent actin spot (AS). Chi-square test: **** *p* < 0.0001. n(AS with indentation) = 93 nuclei, n(AS with no indentation) = 51 nuclei, n(no AS with indentation) = 23 nuclei, n(no AS with no indentation) = 200 nuclei. **d.** Superresolution Airyscan image of a single basal plane of an acinus with an actin spot near the center of the basal surface (white arrowheads). The green dotted box marks the region used to generate orthogonal views (right panel) showing the actin spot (white arrowheads) and adjacent nucleus. Cells were stained as indicated. Scale bar 10 µm. **e.** Quantification of nuclear positioning relative to the basal surface in cells with an actin spot (AS) or without. Wilcoxon test: **** *p* < 0.0001. n(AS contact) = 110 acini, n(no AS) = 82 acini. **f.** Time-lapse movie showing the dynamics of an actin spot (white arrowheads). Cells were transfected with Lifeact-EGFP to label F-actin and iRFP histone-2B (H2B) to label nuclei. n = 31 actin spots. Scale bar, 2 µm. **g.** Images showing nuclear displacement over a 30 s interval. White arrowheads indicate an actin spot. Outlines: red, nuclear position at time 0; green, nuclear position after 30 s. Scale bar, 4 µm. **h.** Box plot showing nuclear mobility in relation to the presence of an actin spot (AS). Wilcoxon test: ** *p* = 0.0017. n(AS) = 16 nuclei, n(no AS) = 28 nuclei.

The indentation of the nucleus around the basal actin spot suggested a physical anchoring of the nucleus. Consistent with this, quantification of nuclear positioning relative to the basal surface showed that nuclei with an adjacent actin spot were significantly closer to the basal membrane compared to nuclei without, which were more evenly distributed along the apico-basal axis (Figure. 3e).

The lack of an actin spot in every cell, suggested the possibility that it was a transient structure. To test this, we generated confocal movies of acini expressing Lifeact-EGFP to label F-actin and H2B-iRFP to label nuclei. Analysis of the movies showed that the basal actin spot was dynamic forming near a basally localized nucleus and persisting for 35 min on average (Figure. 3f). These movies also provided an opportunity to further test the possibility that the actin spot was anchoring the nucleus. We tracked the outline of nuclei every 10 s over 30 min periods. Nuclei associated with a basal actin spot displayed significantly reduced displacement compared to nuclei that were not associated with an actin spot (Figure. 3g–h). These data support the conclusion that the actin spot forms in close association with the nucleus and that the presence of an actin spot changes nuclear shape and mobility, suggesting a mechanical interaction.

### The LINC complex is required to connect the nucleus to the actin spot

To test the dependency of the actin spot on the LINC complex and the nucleus, we quantified the number of actin spots in 3D cultures (day 7) and assessed nuclear positioning in both control and nesprin-2G-depleted cells. The percentage of cells possessing an actin spot was substantially reduced in nesprin-2G knockdown cells compared to controls (Figure 4a,b). Nuclear positioning was also disrupted in the absence of nesprin-2G (Figure 4c). Since acinar tissue architecture was strongly affected by nesprin-2G knockdown with the lumen completely filled as opposed to an open lumen in controls, we also examined an earlier stage (day 4) of acinar development before formation of lumens. In this case, the percentage of cells with an actin spot was also reduced in nesprin-2G-depleted cells (Figure 4d).

**Figure 4.**
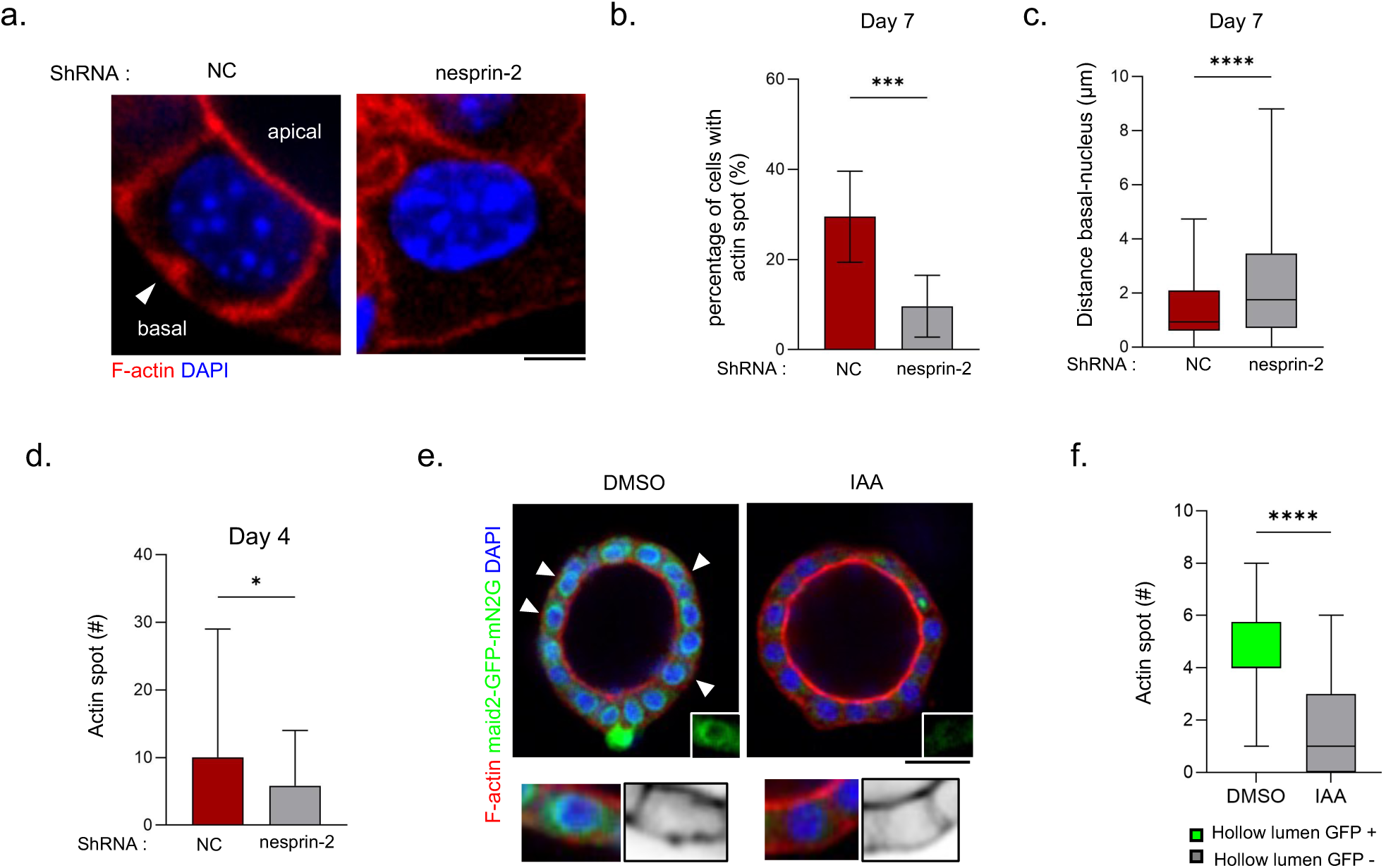
Disruption of the nesprin-2 dirupts the formation of the actin spot. **a.** Representative images of NMuMG acini expressing either noncoding (NC) shRNA or expressing nesprin-2 shRNA at 7 d of development. Note the actin spot (white arrowhead) in the NC shRNA expressign cell. Scale bar, 4 µm. **b.** Histogram showing the percentage of cells with an actin spot in control NC shRNA (red) compared to nesprin2 shRNA (grey) at 7 d development. Unpaired t-test: *** *p* ≤ 0.001. n(NC) = 11 acini, n(nesprin-2) = 9 acini. **c.** Box plot showing nuclear position relative to the basal surface in control NC shRNA (red) compared to nesprin-2 shRNA 2 (grey) at 7 days. Mann-Whitney statistical test shows **** *p* value ≤ 0.0001. n(NC)= 192 nuclei; n(nesprin-2)= 177 nuclei. **d.** Histogram showing the number of actin spot at 4 days in control NC shRNA treated acini(red) vs nesprin-2 shRNA treated (grey). Single confocal planes were used to determine acini numbers. Unpaired t-test: * *p* ≤ 0.05. n(NC) = 22 acini, n(Nesprin-2) = 25 acini. **e.** Representative images showing mAID-EGFP-mN2G in DMSO-treated acini with visible actin spots (white arrowheads), compared to 5-Ph-IAA–treated acini, where both mAID-EGFP-mN2G and actin spots were reduced. Cells were stained as indicated. Scale bar, 30 µm. **f.** Box plot showing the number of actin spot after DMSO or 5-Ph-IAA treatment using the degron system. Only acini expressing EGFP were scored in DMSO treated samples whereas only acini lacking EGFP were scored in IAA treated samples. Mann-Whitney statistical test: **** *p* value ≤ 0.0001. n(DMSO) = 16 acini, n(IAA) = 22 acini.

To test whether the actin spot in fully formed acini was affected by the acute loss of nesprin-2G functionality, we localized actin in the mN2G-degron system. We focused on early times after adding 5-Ph-IAA before lumens had collapsed. As shown in Figure 4e,f, the number of actin spots decreased significantly in 5-Ph-IAA treated acini compared to vehicle controls. Also note that the nuclei in 5-Ph-IAA-treated acini have lost the EGFP signal from the mN2G-degron, confirming that the degron had been activated (Figure 4e).

### The actin spot reflects the formation of a nuclear-plasma membrane contact through the LINC complex and β1-Integrin

The LINC complex can exhibit polarized localization within the nuclear envelope during morphogenetic events. For example, during fibroblast polarization for migration, nesprin-2-SUN2 LINC complexes align along actin cables on the dorsal nuclear surface (*5*). In cells migrating through confined spaces, nesprin-2G accumulates at the front of the nucleus (*38*). During cytomegalovirus infection, SUN1 accumulates on the surface of the nucleus adjacent to the virus assembly compartment (*39*).

Given these findings, we wondered whether the LINC complex might also be polarized when the nucleus is indented and adjacent to the actin spot. In EGFP-SUN2-expressing NMUMG acini, we observed a clear accumulation of SUN2 in the indented nuclear surface facing the actin spot (Figure 5a,b). In acinar cells lacking an actin spot and nuclear indentation, EGFP-SUN2 appeared uniformly distributed around the nuclear envelope (Figure 5a, b). These data point to a physical and functional engagement between the LINC complex and the actin spot.

**Figure 5.**
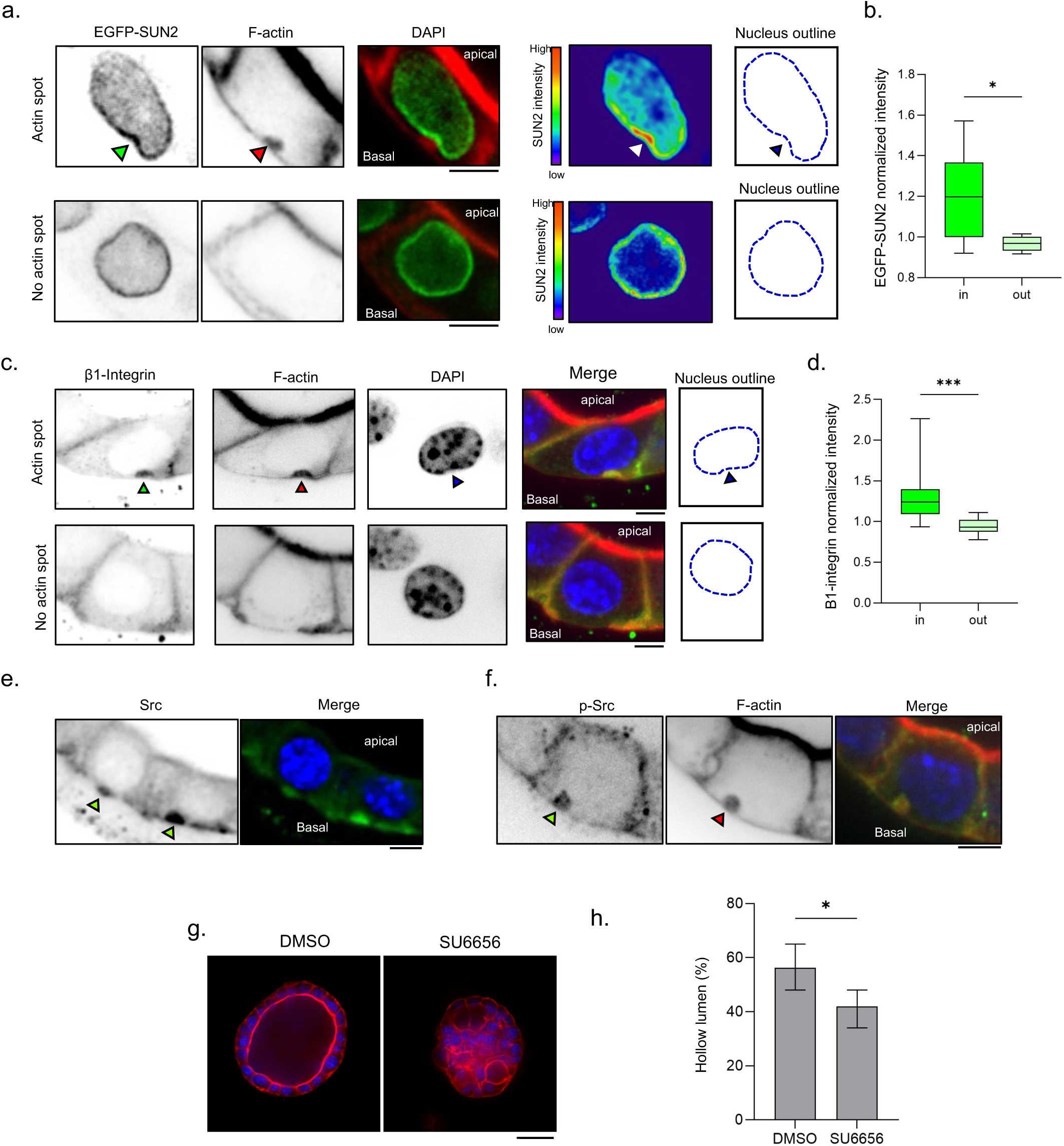
Accumulation of EGFP-SUN2, β1-Integrin and Src near or in the actin spot and requirement of Src activity for maintenance of acini architecture. **a.** Top panel, confocal images showing a nucleus adjacent to a basal actin spot (red arrowhead), exhibiting localized accumulation of EGFP-SUN2 (green arrowhead). The heatmap (Royal LUT, ImageJ) indicates EGFP-SUN2 intensity. The blue outline shows an indented nucleus. Bottom panels, confocal images showing a nucleus without an adjacent actin spot, displaying uniform EGFP-SUN2 distribution around the nuclear envelope (heatmap) and a rounded nucleus (outlined in blue). Scale bar, 4 µm. **b.** Box plot quantifying EGFP-SUN2 intensity adjacent to the actin spot (“in”) compared to the rest of the nuclear envelope (“out”), normalized to the mean EGFP-SUN2 intensity of the entire nuclear envelope. Wilcoxon statistical test: * p value ≤ 0.05. n= 11 nuclei. **c.** Top panels, confocal images showing a nucleus adjacent to an actin spot (red arrowheads), with localized accumulation of β1-Integrin (green arrowheads) at the actin spot. The blue outline shows an indented nucleus. Bottom panels, confocal images showing a nucleus without an adjacent actin spot and lacking β1-integrin accumulation. Scale bars, 4 µm. **d.** Box plot quantifying β1-Integrin intensity at the basal membrane region associated with the actin spot (“in”) compared to the rest of the basal membrane (“out). Wilcoxon statistical test: * p value ≤ 0.05. n =15 cells. **e.** Confocal images of Src localization on the basal surface (green arrowheads) in proximity to the nucleus. Scale bar, 4 µm. **f**. Confocal images showing pSrc (green arrowhead) colocalizing with a basal actin spot (red arrowhead). Scale bar, 4 µm. **g**. Conmfocal images of a vehicle (DMSO) treated acini with a hollow lumen compare to a collapse lumen after treatment with 10µM of SU6656 for 20 h. Scale bar, 30 µm. **h.** Histogram showing the percentage of hollow acini in DMSO vs SU6656 treatments. Unpaired t-test: ** *p* value ≤ 0.01. n(DMSO) = 1022 acini, n(SU6656) = 817.

Since the actin spot was localized on the basal side of the epithelium, we next determined whether other basal epithelial components might be colocalized with it. Integrins are key components of the basal surface of epithelia and inhibition of β1-integrin is associated with defects in lumen formation (*20, 40, 41*). As expected, β1-integrin was mainly localized to the basolateral domain of luminal cells (Figure 5c,d). In cells possessing an actin spot, we observed a distinct cluster of β1-integrin that colocalized with the actin spot and was adjacent to the indented nucleus (Figure 5c-d). Given that β1-integrin is a membrane protein, this indicates that the actin spot reflects F-actin localized on an indented plasma membrane. Combined, the localization data support the existence of a nuclear–plasma membrane contact site mediated by the LINC complex, F-actin and β1-integrin.

As other membrane contact sites act as signaling platforms, we hypothesized that the nuclear-plasma membrane contact site might contain signaling components. We focused on integrin associated molecules such as those found in focal adhesions. We did not observe talin or FAK in the contact site, but we did observe a striking accumulation of Src adjacent to i nuclei (Figure 5e). A second Src antibody, this one to active pSrc, also decorated the actin spot (Figure 5f). To test whether Src activity played role in maintaining acinar architecture, we treated acini with the Src inhibitor, SU6656. Treatment of acini with 10 µM SU6656 for 20 h reduced the number of acini with hollow lumens (Figure 5g,h). Combined, these data suggest that the nuclear-plasma membrane contact may activate Src to control epithelial architecture.

## Discussion

Functions of the LINC complex in epithelial tissues and the consequences of its downregulation such as occurs in carcinomas are poorly understood. Our study highlights a role for nesprin-2G via its actin-binding domain and interaction with FHOD1 in mammary gland morphogenesis and maintaining acinar architecture. We show that the nucleus is transiently positioned and anchored at the basal surface of the cell by the local accumulation of a basal actin spot correlated with the presence of the LINC complex, β1-integrin, and active Src. We show that degradation of nesprin-2G and inhibition of Src both lead to acini with collapsed lumens. Coupled with earlier studies showing similar acinar collapse when β1-integrin is inhibited or Rho is elevated (*20, 40*), our data suggest a coordinated mechanochemical signaling pathway in which the proximity between the basal membrane and the nucleus activates a β1-integrin-Src pathway to sustain homeostatic contractility in the acinar epithelium (Figure 6).

**Figure 6.**
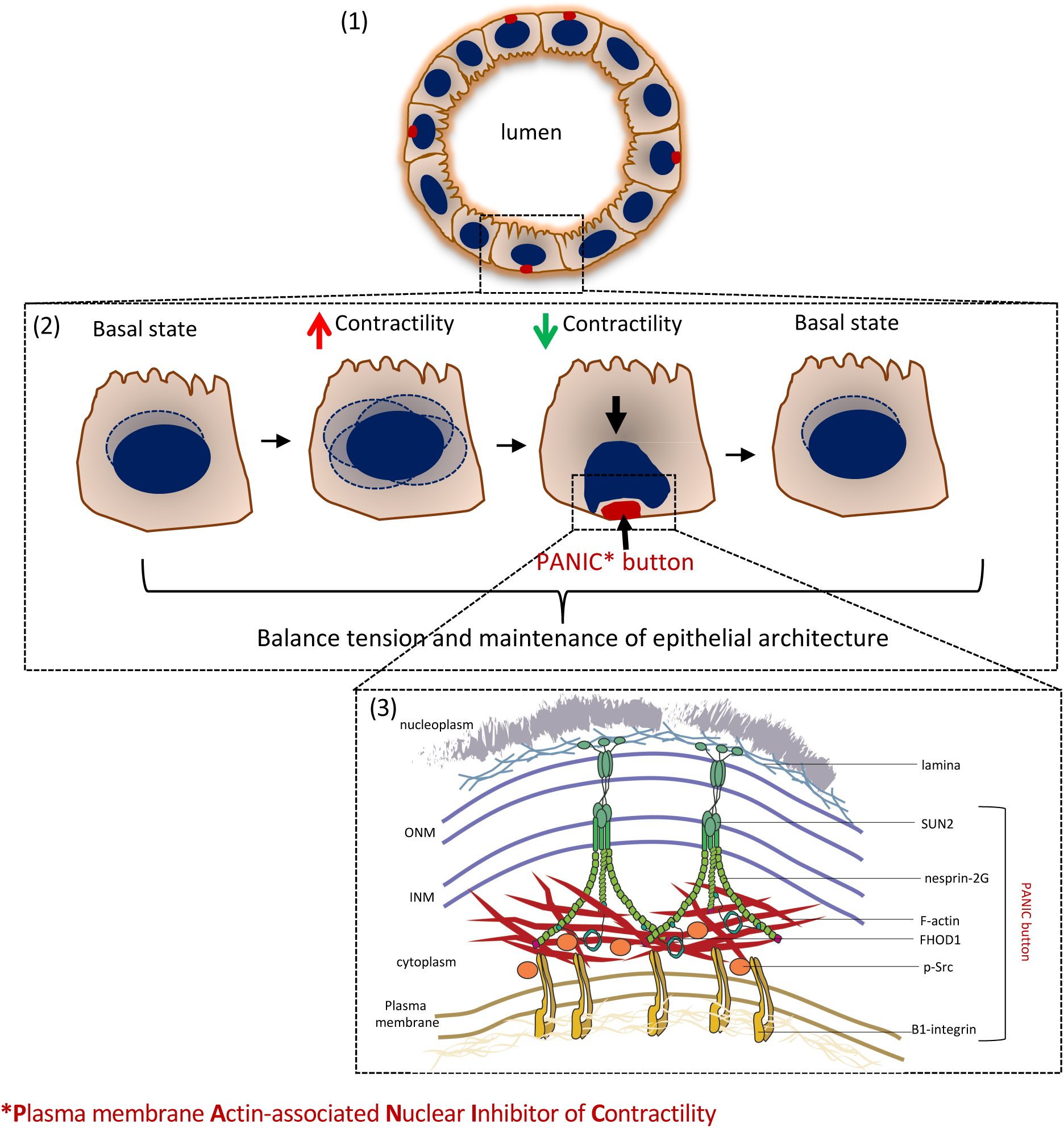
The PANIC button working model for homeostatic regulation of epithelial architecture by the nucleus. (1) The maintenance of a hollow lumen in glandular epithelial acini requires tight control of cell contractility. (2) When cellular contractility increases, this triggers nuclear displacement. If contractility is sufficient to displace the nucleus to the basal surface, it generates the nuclear-plasma membrane contact site or PANIC button. (3) The PANIC button is signaling hub composed of SUN2, nesprin-2G, FHOD1, β1-Integrin and Src. We propose that this signaling hub initiates a negative feedback loop through Src signaling that reduces cell contractility and restores tissue homeostasis.

It is well established that mechanical signals originating at the cell surface — particularly through integrins — can be transmitted to the nucleus via the cytoskeleton and the LINC complex (*42, 43*). However, to our knowledge, it has never been proposed that the nucleus might form a direct or spatially restricted contact with the plasma membrane in response to mechanical stimuli. Membrane contact sites (MCS) are defined as regions where the membranes of two organelles are closely apposed — typically within 10–50 nm — without fusing, and where specific tethering proteins enable functional communication, such as lipid exchange or calcium signaling. Well-characterized MCS include ER–mitochondria, ER– plasma membrane, and ER–endosome contacts, which are now recognized as key hubs of intracellular organization and signaling (*44, 45*).

While membrane contact sites between intracellular organelles are well described, a functional interface between the plasma membrane and the nuclear envelope has not been identified previously. In our study, we observe the accumulation of β1-integrin and Src in close proximity to the basal nuclear envelope where SUN2 accumulates, suggesting a subcellular arrangement consistent with a membrane contact site. Although ultrastructural analysis will be required to definitively confirm the presence of a physical membrane apposition, our data support the idea that this proximity — stabilized by the LINC complex connections to F-actin — could act as a signaling axis for Src activation and be essential for maintaining acinar homeostasis.

How does the pathway we have identified regulate acinar architecture? We propose that the LINC complex, F-actin, β1-integrin and Src pathway regulates Rho levels and cellular contractility to prevent acinar collapse. Previous work has shown that all of the factors in this pathway regulate Rho stimulated contractility in glandular epithelia and that activation of Rho is sufficient to induce acinar collapse (*19, 20*). Src is known to regulate two Rho GTPase pathways at focal adhesions: An early pathway at clustering integrins in which Src phosphorylates and activates p190 RhoGAP to reduce Rho GTP levels (*46*) and a later pathway at maturing focal adhesions in which Src phosphorylates and activates the Rho GEF LARG to increase Rho GTP levels (*47*). We propose that the nuclear-plasma membrane site controls Rho levels via the “early” integrin signaling pathway. Consistent with this idea, p190RhoGAP is critical for ductal morphogenesis of the mammary gland during development (*48, 49*).

Combining this molecular pathway with the specific nuclear-membrane contact site, leads to a model in which the nucleus acts as a mechanical sensor of cellular contractility: when Rho GTP levels are in the normal range the nucleus remains near the cell centroid, but when contractility becomes elevated, nuclear displacement is elevated leading to the formation of the nucleus-plasma membrane contact site and the signaling pathway that reduced Rho GTP levels and cellular contractility. In this scenario, the nucleus plays the role of a mechanical sentinel of cellular contractility. Given this function, we think an appropriate name for the nuclear-plasma membrane contact site is: <u>P</u>lasma membrane-<u>A</u>ssociated <u>N</u>uclear Inhibitor of <u>C</u>ontractility (PANIC) button. It will be interesting to explore how loss of the PANIC button, such as would be predicted to occur in cases where the LINC complex is down regulated, (e.g., in breast cancer and other cancinomas (*25–27*), contributes to the development of disease.

## Material and Methods

The sources of all reagents used are indicated in the Supplementary Table following the Supplementary Figure Legends.

### Culture cell

All cells were maintained at 37 °C in a humidified incubator with 5% CO₂. NMuMG (Normal Murine Mammary Gland) cells were from ATCC and were cultured in Dulbecco’s Modified Eagle Medium (DMEM) supplemented with 10% fetal bovine serum (FBS), 1X penicillin-streptomycin mix (Pen-Strep), and 10 µg/ml insulin. MCF-10A human breast epithelial cells were obtained from ATCC and maintained as previously described (*25*). Culture of cells to generate 3D acini followed a protocol adapted from the Brugge laboratory (*25*). Forty-five µL of Matrigel (Corning) was added to each well of an 8-well chamber slide (™ Lab-Tek™ Chamber Slide Thermo Fisher or µ-Slide 8 Well Glass Bottom/polymer, Ibidi). A suspension of cells was prepared in Assay Medium (DMEM/F12 supplemented with 0.5 mg/ml hydrocortisone, 100 ng/ml cholera toxin, 100 μg/ml insulin, 1X Pen-Strep and 2% (v/v) horse serum) containing 5 ng/ml epidermal growth factor (EGF) and 2% Matrigel. Each well was seeded with 5000 cells. Cultures were fed every four days with fresh Assay Medium containing 2% Matrigel and 5 ng/ml EGF. Acini with hollow lumens developed in 7 days for NMuMG cells and 15 days for MCF-10A cells.

### Plasmids

All constructs were confirmed by DNA sequencing. pMSCV-puro EGFP-C4 retroviral vector was previously described (*31*). It was used to make retroviral particles in 293T cells and to express N-terminal EGFP fusion protein in cells. pLKO2-hygro H1 lentiviral vector was derived from TRC2-pLKO-puro (Millipore-Sigma, SHC201) plasmid by replacing the puromycin selectable marker gene and human U6 promotor with a hygromycin selectable marker gene from pMSCV-hygro (Clontech) and human H1 promoter from pSUPER.retro.puro (Oligoengine), respectively. The resultant plasmid was used to make lentiviral particles in 293T cells and to express shRNA in mammalian cells.

Constructs (with insert restrictions sites) for Lifeact (BamHI/NotI), miniN2G (NotI), hFHOD1 WT (BamHI/NotI), hFHOD1 R136A R137A (BamHI/NotI), and hSUN2 WT (BglII/NotI) were previously described (*29, 34*). The shRNA sequences for NC (Sigma, 5’-CAACAAGATGAAGAGCACCAA-3’), mFHOD1-4 (Sigma, TRCN0000216791, 5’-GAACCTCTTTCCTACCATTTC-3’), hSUN1-1 (Sigma, TRCN0000297311, 5’-GCTGTTCTGAAACTTACGAAA-3’), hSUN2-1 (Sigma, TRCN0000143335, 5’-GCCTATTCAGACGTTTCACTT-3’), respectively. The shRNA sequences for hNesprin-1, hNesprin-2, mNesprein-2, and mFHOD1 were previously published (*30, 31, 50*). pLVX EF1a H2B-iRFP670 IRES-blast was obtained from Dr. Hee-Won Yang (Columbia University).

### Transfection, viral infection, and selection

For production of viruses, 293T cells were transfected using the calcium phosphate method and viruses were harvested as previously described (*31*). NMuMG and MCF10A cells were infected with viral supernatant as previously described (*31*).

### Description of the Degron system

We employed the auxin-inducible degron2 system to generate NMuMG cells that would allow rapid inactivation of nesprin-2G actin functionality. First, NMuMG cells were stably knocked down for nesprin-2G using shRNA as above. Next, we transfected the nesprin-2G depleted cells with a mAID2-EGFP-mN2G degron construct that contains the auxin-inducible degron2 tag (*36*), a EGFP tag and mini-nesprin2G, which contains the actin binding functionality of nesprin-2G attached to the KASH domain (*5*). Expressing cells were then transfected with the degradation enzyme OsTIR1(F74G). As shown in Figure 2, the endogenous nesprin-2G protein was effectively silenced by the shRNA, the miniN2G-degron was expressed, localized properly to the nucleus and rescued lumen formation in knockdown cells. We used the auxin 5-Ph-IAA at 200 nM for 2D cells and 5 µM 3D cells to induce degradation.

### Drug treatments

To activate the inducible-auxin degron system, 2D monolayers of NMuMG cells were treated with 400 nM 5-Ph-IAA for 30 min to 2 h, followed by fixation or live-cell imaging. For 3D cultures, wild-type NMuMG cell acini with hollow lumens (day 7 of culture) were incubated with 5 µM 5-Ph-IAA for 24 h prior to fixation with 4% paraformaldehyde. For live imaging, samples were imaged for a minimum of 15 h.

NMuMG cell acini (7 days of culture) were treated with Src family kinase inhibitor 10 µM SU6656 for 20 h. Following treatment, samples were directly imaged in bright field or fixed with 4% paraformaldehyde and stained with phalloidin to visualize F-actin.

### Western blotting

Cells were washed with PBS and lysed on ice using 1X Laemmli SDS sample buffer. Lysates were collected by scraping, transferred to tubes on ice, then boiled for 5 min and sonicated. Protein samples were subjected to SDS-PAGE. Proteins were transferred onto nitrocellulose membranes (Cytiva), blocked with non-fat dry milk (LabScientific) and then incubated with primary and secondary antibodies. The signal on the membrane was detected by LI-COR Odyssey Imaging System.

Primary antibodies (see Supplementary Table for sources) used for blotting included anti-nesprin2G (rabbit, 1:2000); anti-nesprin 1 (mouse, 1:500); anti-SUN1 (rabbit, 1:2000); anti-SUN2 (rabbit, 1:2000); anti-tubulin (rat, 1:2000); anti-FHOD1 (rabbit, 1:2000).

### Immunostaining

2D monolayer cells were fixed with 4% paraformaldehyde in PBS for 15 minutes, permeabilized with 0.2–0.5% Triton X-100 in PBS and then blocked 3–5% BSA in 0.2% Triton X-100. The samples were then incubated with primary and fluorophore-conjugated secondary antibodies and fluorophore-conjugated phalloidin. Coverslips were mounted using Fluoromount-G with DAPI.

Acini were immunostained following an established protocol from the Brugge laboratory (*25*). Briefly, the acini were fixed in 4% paraformaldehyde for 20 minutes, permeabilized, and blocked before incubation with primary and secondary antibodies and with phalloidin. Finally, the acini are covered by Fluorochrome-G mounting media in Ibidi 8-chamber polymer or glass bottom.

Primary antibodies (see Supplementary Table for sources) used for immunofluorescence include: anti-nesprin2G (rabbit, 1:100); β1-integrin (rabbit, 1:100); Src (mouse, 1:100), pSrc (Tyr416) (rabbit, 1:100). Secondary antibodies were Alexa Fluor 488 (rabbit, mouse, 1:100); Alexa Fluor 568 (rabbit, mouse, 1:100), Alexa Fluor 647 (rabbit, mouse, 1:100);

### Microscopy and live imaging

Images of 2D monolayer cells were acquired with 60× Apo TIRF oil (NA 1.49) objective lens and a Nikon DS-Qi2 CMOS CCD camera on an inverted Eclipse Ti microscope (Nikon) equipped with SPECTRA X Light ENGINE (Lumencor, Inc.) light source controlled by NIS Elements software (Nikon). Confocal images of 3D acini were acquired with a Nikon A1R MP confocal scanner mounted on a TiE inverted microscope with 40x PlanFluor oil (NA 1.3) objective lens. The microscope was controlled by NIS Elements software. Alternative confocal images were obtained with a X-Light V3 Spinning Disk (CrestOptics) mounted on inverted Ti2 microscope with 10x, 20x 40x PlanApo air (NA 0.3, 0.8, 0.95, respectively) objective lens and Hamamatsu ORCA-Fusion BT CMOS CCD camera. Super-resolution images were acquired using an LSM 900 Airyscan confocal scanner mounted on an AxioObserver 7 inverted microscope (Zeiss) with a 63x PlanApo oil (NA 1.4) objective lens. Live imaging was performed by 40x and 60x PlanApo oil objective (NA 1.3 and 1.45, respectively) using Andor BC43 microscope (Oxford instruments), and by Andor Fusion software.

### Image analysis

#### Actin spot dynamics

We generated movies of NMUMG 3D acini expressing Lifeact-EGFP (to probe actin) and H2B-iRFP (to probe nuclei) using scanning confocal microscopy. Images were acquired on a single plane every 10 min to allow the lifetime of the actin spot to be determined. We used IMARIS software to analyze the dynamics of the actin spot.

#### Nuclear tracking

From movies as above for actin spot dynamics, we compared the mobility of nuclei every 10 sec over a 30 min time course. We compared mobility of nuclei in contact with the actin spot to those that were not contacting the actin spot. We measured nuclear stability by creating surfaces or spots on the nucleus using IMARIS software to collect their displacement over time. We also created a mask on nuclei on ImageJ to show superposed to different time and highlight the nuclear displacement.

### Quantification and statistics

Statistical analyses were performed with Prism software. The number of replicates (N) and p-values are indicated in the figure legends and in the data tables. Box plots were generated in Prism and represent the median of 10 and 90 percentile.

### Lumen quantification

Acini were visually inspected and scored as containing a hollow lumen or a filled (non-hollow) lumen after taking images to the microscope. from three independent experiments. The proportion of hollow acini is presented as a percentage in the figures. A two-tailed unpaired t-test was applied to compare two different groups, and a one-way analysis of variance (ANOVA) followed by Tukey’s post hoc test was used for multiple group comparisons.

### Nuclear positioning and mobility

Nuclear positioning relative to the basal surface was determined by measuring the distance from the basal surface to the basal edge of the nucleus using ImageJ software or IMARIS. The difference in nuclear positioning between a nucleus in or non-in proximity with an actin spot was statistically tested by a Wilcoxon test (paired – non parametric samples). The difference in nuclear positioning between control and nesprin-2G knockdown condition was tested by Mann-Whitney (unpaired – non parametric samples) statistical test.

Nuclear mobility was quantified using IMARIS software with automated spot tracking. The speed (μm/sec) of each nucleus was measured and normalized to the mean speed of nuclei that were not in proximity to an actin spot. To assess whether nuclear mobility was influenced by proximity to actin, the difference in mobility between nuclei in proximity to an actin spot and those not in proximity was statistically evaluated using a Wilcoxon test (paired, nonparametric samples).

### EGFP-SUN2 and β1-integrin intensity

For the measurements in Figure 5a-d. The intensity of Integrin-β1 and EGFP-SUN2 was measured in ImageJ by drawing a line along the nuclear surface (EGFP-SUN2) or plasma membrane surface (β1-integrin) directly adjacent to the actin spot and another line along the surfaces not adjacent to the actin spot. The intensities were then normalized to the total intensity of the entire nuclear surface for EGFP-SUN2 or the entire basal cell surface for β1-integrin. The difference in the fluorescence intensities in or not in contact with actin spot statistically tested by a Wilcoxon test (paired – non parametric samples).

## Acknowledgements

We thank Theresa Swayne in the Confocal and Specialized Microscopy core (Herbert Irving Comprehensive Cancer Center, Columbia University) for help with microscopy and access to their Zeiss Airy Scan confocal microscope and Nikon scanning confocal. We thank Rivka Shapiro for the images analysis. This work was supported by NIH grants R35 GM136403 to GGG and UO1 CA2255663 to TL. The content is solely the responsibility of the authors and does not necessarily represent the official views of the NIH.

## Supplementary Figure Legends

**Supplementary Figure 1. Knockdown of specific LINC complex proteins and FHOD1. a**. Western blot analysis of lysates from stable MCF-10A cell lines expressing shRNAs targeting individual LINC complex genes, confirming effective knockdown. **b.** Immunofluorescence staining of stable NMuMG cell lines expressing nesprin-2G shRNA, confirming nesprin-2 knockdown. **c.** Western blot showing knockdown efficiency of FHOD1 shRNA, compared to control shRNA in NMuMG cell lysates. Scale bar: 20 µm

**Supplementary Figure 2. Degron-system depletion of Nesprin-2G and validation. a.** Images of a 2D monolayer showing the time-dependent disappearance of mini-N2G-EGFP in response to 5-Ph-IAA (200nM) treatment compared to DMSO control. Scale bar: 20µm. **b.** Quantification of the percentage of spheroids with hollow lumens in wild-type NMuMG cells treated with DMSO or 5-Ph-IAA (5µM). Statistical significance was assessed using chi-square test, ns is for non-significant.

## Supplementary Table (following pages)

Supplementary Table (following pages):
| items | source | Identifier |
| --- | --- | --- |
| **Antibodies** |  |  |
| Rabbit polyclonal anti-nesprin-2 | (5) | S2-CH |
| Mouse monoclonal anti-nesprin-2 | Wolfson CIND | MANNES2G(4B5) |
| Mouse monoclonal anti-nesprin-1 | Wolfson CIND | MANNES1E |
| Rabbit polyclonal anti-SUN1 | SIGMA | HPA008346 - 100 uL |
| Rabbit polyclonal anti-SUN2 | abcam | ab87036 |
| Rabbit polyclonal FHOD1 | Santa Cruz Biotech | sc-99209 |
| Rabbit polyclonal anti-integrin-b1 | Thermofisher | PA5-78028 |
| Mouse monoclonal anti-src | Thermofisher | MA5-11173 |
| Rat polyclonal tyrosinated alpha-tubulin | European Collection of Authenticated Cell Cultures | 92092402 |
| Alexa fluor 568 anti-rabbit | Jackson ImmunoResearch Laboratories | A10042 |
| Alexa fluor 568 anti-mouse | Jackson ImmunoResearch Laboratories | A10037 |
| Alexa fluor 488 anti-rabbit | Jackson ImmunoResearch Laboratories | 711-545-152 |
| Alexa fluor 488 anti-mouse | Jackson ImmunoResearch Laboratories | 715-545-150, |
| Alexa fluor 647 anti-rabbit | Jackson ImmunoResearch Laboratories | 711-605-152 |
| Alexa fluor 647 anti-mouse | Jackson ImmunoResearch Laboratories | 715-605-151 |
| IRDye 680 conjugated goat anti-mouse IgG | LI-COR Biosciences | 715-605-151 |
| IRDye 680 conjugated goat anti-rabbit IgG | LI-COR Biosciences | 926-32221, |
| IRDye 680 conjugated goat anti-rat IgG | LI-COR Biosciences | 926-32229, |
| IRDye 800CW conjugated goat anti-mouse IgG | LI-COR Biosciences | 827-08364 |
| IRDye 800CW conjugated goat anti-rabbit IgG | LI-COR Biosciences | 925-32219, |
| IRDye 800CW conjugated goat anti-rat IgG | LI-COR Biosciences | 926-32219 |
| **Recombinant DNA** |  |  |
| pMSCV-puro EGFP-C4 | (31) |  |
| pMSCV-puro EGFP-C4 miniN2G | (29) |  |
| pMSCV-puro EGFP-N4 Lifeact | (29) |  |
| pMSCV-puro EGFP-C4 hFHOD1 WT | (31) |  |
| pMSCV-puro EGFP-C4 hFHOD1 R136A R137A | (34) |  |
| pMSCV-puro EGFP-C4 hSUN2 WT | This paper |  |
| pLVX EF1a H2B-iRFP IRES-Blast | Hee-Won Yang |  |
| pSUPER.retro.puro NC | (31) |  |
| pSUPRE.retro.puro hNesp1-1 | This paper |  |
| pSUPRE.retro.puro hNesp2-1 | This paper |  |
| pSUPRE.retro.puro hSUN1-1 | This paper |  |
| pSUPRE.retro.puro hSUN2-1 | This paper |  |
| pLKO2-hygro hH1 NC | This paper |  |
| pLKO2-hygro hH1 mNesp2-2 | This paper |  |
| pLKO2-hygro hH1 mFHOD1-1 | This paper |  |
| pLKO2-hygro hH1 mFHOD1-4 | This paper |  |
| pHIT 60 | (51) |  |
| psPAX2 | Addgene | 12260 |
| pMD2.G | Addgene | 12259 |
| **Chemical reagent** |  |  |
| Rhodamin-Phalloidin | ThermoFisher | R415 |
| Alexa Fluor 488-phalloidin | Thermo Fisher Scientific | A12379 |
| Alexa Fluor 647-phalloidin | Thermo Fisher Scientific | A22287 |
| 5-ph-iaa | MedChemExpress (MCE) | HY-134653 |
| CellMask™ Plasma Membrane Stains | C10046 |  |
| Sir-DNA 555 |  |  |
| Matrigel (Matrix basement membrane) | corning | 356231 |
| PP2 | Sigma | 529576 |
| SU6656 | MedChemExpress (MCE) | HY-B0789 |
| Paraformaldehyde | Fisher | 50-980-494 |
| Items | Source | identifier |
| **Experimental model: cell lines** |  |  |
| NMuMG | ATCC | CRL-10317 |
| MCF10A | ATCC | CRL-1636 |
| **Software and Algorithms** |  |  |
| Fiji | ImageJ |  |
| IMARIS | Oxford instruments |  |
| PRISM | GraphPad |  |

## Notes

### Competing Interest Statement

The authors have declared no competing interest.

