## Supplementary figures and images for "The nucleus forms a dynamic contact with the plasma membrane to maintain the glandular epithelial architecture"

### supplemental Figures

#### Supplementary Figure 1

Rayer et al. 2025

a.

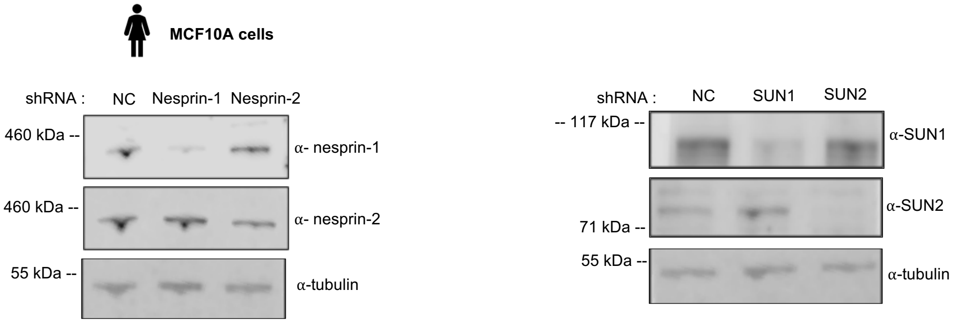


b.

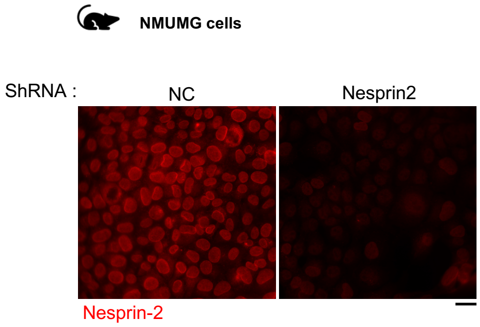


c.

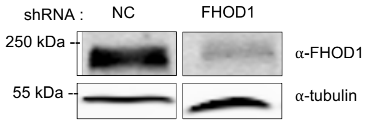

#### Supplementary Figure 2



a.

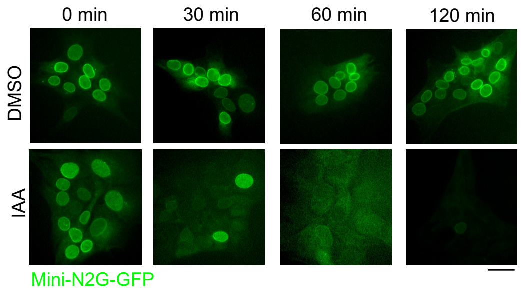


b.

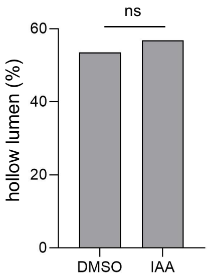
